# American ginseng (*Panax quinquefolius* L.) extracts (G1899) reverse stress-induced behavioral abnormalities in mice

**DOI:** 10.1101/2025.03.21.644575

**Authors:** Rahmi Lee, McKennon J. Wiles, Ellison R. Black, Seung Hyun Roh, Evelina Bouckova, Madison H. Wustrau, Joshua C. Flowers, Paige E. Vetter, Jaehoon Lee, Byung-Cheol Han, Seonil Kim

## Abstract

Stress affects brain functions, which leads to the development of mental disorders like anxiety, depression, cognitive decline, and social dysfunction. There is increasing focus on the role of nutritional, herbal and nutraceutical compounds on mental and cognitive functioning. Interestingly, studies suggest that American ginseng (*Panax quinquefolius* L.) extracts (G1899) improve cognition. We thus examined whether G1899 showed protective effects on stress-induced behavioral changes in animals. 200 mg/kg G1899 was orally administered daily for 4 weeks to 2-3-month-old female and male mice before inducing stress. To induce acute stress in animals, we intraperitoneally injected a low dose of lipopolysaccharides (LPS) (10 μg/kg), and saline was used as a control. We also used chronic restraint stress (CRS) as a chronic stress model in mice. After LPS injection or CRS, multiple behavioral assays were carried out – a sucrose preference test, an open filed test, reciprocal social interaction, contextual fear conditioning, and a tail suspension test – to determine whether acute or chronic stress affected animals’ behaviors and whether G1899 had protective effects against stress-induced behavioral dysfunction. We found that both LPS injection and CRS induced stress-related behavioral dysfunction, including depression-like behavior, anhedonia, social dysfunction, and fear memory impairments in both females and males. However, G1899 treatment was sufficient to reverse stress-induced behavioral abnormalities in animals. Our data further suggested that G1899 reduced the activity of hippocampal neurons by suppressing glutamatergic activity. Our study suggests that G1899 supplements can be protective against both acute and chronic stress in mice by suppressing neuronal and synaptic activity.

**Highlights:**

- American ginseng (*Panax quinquefolius* L.) extract (G1899) treatment reverses acute stress-induced behavioral dysfunction in mice.
- G1899 treatment reverses chronic stress-induced behavioral dysfunction in mice.
- G1899 treatment reduces serum corticosterone levels in chronically stressed mice.
- G1899 treatment suppresses glutamatergic activity in hippocampal neurons.

## Introduction

Ginseng is derived from the roots of *Panax ginseng* C.A. Meyer and traditionally used as a medicinal herb in Korea, Japan, China, and the United States (Attele et al., 1999; Choi et al., 1995). Ginseng has utilized in medicine since ancient times because it is suggested to induce anticancer, antioxidative, anti-aging, anti-stress, anti-inflammatory, anti-fatigue, anti-diabetic, antidepressant, and neuroprotective effects (Sana et al., 2024). Importantly, several previous studies suggest that ginseng can affect various functions of the central nervous system including improved blood flow, protected neurons from damage, enhanced neural development, improved learning and memory, reduced neuroinflammation, oxidative stress, and reduced neurodegeneration (Chan et al., 2003; Chang et al., 2003; Helms, 2004; Hou et al., 2020; Kim et al., 2023; Kim et al., 2024; Papapetropoulos, 2007; Scott et al., 2001; Shin et al., 2021; Stavro et al., 2005; Xie et al., 2005). Ginseng’s active constituents, such as saponin in a form called ginsenosides, are thought to be responsible for these effects (Ong et al., 2015). Recent studies also demonstrate that non-saponin fraction with rich polysaccharide from Korea red ginseng induces neuroprotective effects in Alzheimer’s disease model mice (Kim et al., 2023; Kim et al., 2024). Extensive research over the years has shown that ginseng has potential as neuroprotective compounds, which can be used to treat and prevent neurological damage or pathologically related diseases. Among several ginseng species, the root of *Panax quinquefolius* L., or American ginseng, is regarded as a premium dietary supplement and nutritious food that has been used for decades as an edible food and beverage component in America (Qu et al., 2009; Riaz et al., 2019). However, there is currently limited knowledge of the *in vivo* beneficial effects of American ginseng.

According to a survey conducted by the American Psychological Association (APA) in 2021, more than 70% of Americans that experience high stress have various mental problems. These alarming statistics about stress demonstrate the widespread prevalence of this state of mind. Stress has a negative impact on health when it rises above a particular threshold. Repeated stress affects brain functions, contributing to the development of anxiety, depression, cognitive decline, and social dysfunction (daSilva et al., 2021; Kessler, 1997; Lupien et al., 2009). Unfortunately, there is no appropriate treatment for these mental disorders. Interestingly, recent studies have suggested that ginseng is involved in adjusting the hypothalamic–pituitary– adrenal (HPA) axis to produce beneficial effects on the heart and brain (Lee and Rhee, 2017). However, the mechanism underlying the effects of ginseng on these stress-related diseases has not been completely established.

Ginseng is shown to either enhance or suppress neuronal activity, which is presumed to be cellular mechanisms of its neuroprotective effects (Lee et al., 2000; Leem et al., 2020; Nah, 2014; Shin et al., 2012; Wang et al., 2011). In fact, ginsenosides are shown to work by inhibiting the elevation of cytosolic Ca^2+^ and Na+, and by inhibiting glutamatergic NMDA receptors (NMDARs) (Kim et al., 2002; Kim et al., 2004). Interestingly, ketamine is a noncompetitive NMDAR antagonist that inhibits excitatory synaptic transmission (Anis et al., 1983). Ketamine at low doses works as a rapid-acting antidepressant and improves cognitive functions in humans (Keilp et al., 2021; Miller et al., 2016). This ketamine’s actions are likely mediated by NMDAR antagonism (Aleksandrova et al., 2017). After demonstrating rapid and robust antidepressant efficacy, the US Food and Drug Administration (FDA) approved esketamine (the S enantiomer form of ketamine) for the treatment of depression in 2019 (Kohtala, 2021). Additionally, several studies including our own suggest that ketamine may have neuroprotective effects against chronic stress-related neuropsychiatric conditions in addition to decreasing depression in both humans and animals (Brachman et al., 2016; Feder et al., 2014; Flowers et al., 2024; Gill et al., 2021; Lee et al., 2021b; Ma et al., 2019; Ma et al., 2022; Taylor et al., 2018; Yang et al., 2018). Therefore, NMDAR antagonism by ginseng may underlie its protective effects on brain disorders.

Here, we discover that American ginseng (*Panax quinquefolius* L.) extracts show protective effects on acute and chronic stress-induced behavioral dysfunction in both female and male mice. We also find that G1899 significantly decreases the activity in hippocampal neurons by reducing glutamatergic activity. Our study thus suggests that American ginseng supplements can be protective against both acute and chronic stress in animals by suppressing neuronal and synaptic activity.

## Materials and methods

### Animals

C57Bl6J (Jax 000664) and CD1 (ICR) (032) mice were obtained from Jackson laboratory and Charles River, respectively, and bred in the animal facility at Colorado State University (CSU). Animals were housed under a 12:12 hour light/dark cycle. CSU’s Institutional Animal Care and Use Committee (IACUC) reviewed and approved the animal care and protocol (3408). As no sex difference was found in our experiments, we presented data combined from male and female mice.

### American ginseng (*Panax quinquefolius* L.) extract (G1899) manufacturing

The processed *Panax quinquefolius* was meticulously cleaned to eliminate any foreign materials, finely chopped, and extracted using a water-to-material ratio of 9:1 at 90°C. The obtained extract was subsequently dried via spray-drying, achieving a final moisture content of 4% (±1%). The resulting G1899 extract was then resuspended in a suitable solvent for further analytical and experimental investigations.

### G1899 Preprocessing for Analysis

0.5 g of G1899 was resuspended in 10 mL of 70% MeOH and subjected to sonication for 30 min. Following sonication, the sample was centrifuged at 5000 × *g* for 10 minutes, and the supernatant was collected. The obtained supernatant was then filtered through a 0.2 μm PVDF membrane filter. The final extract was appropriately diluted and utilized for ultra-performance liquid chromatography (UPLC) and ultra-high performance liquid chromatography-tandem mass spectrometry (UHPLC-MS/MS) analyses.

### UPLC

Chromatographic analysis was performed on a Waters ACQUITY UPLC equipped with on ACQUITY UPLC BEH Phenyl column (2.1 × 10 mm, 1.7 μm). The column temperature was 35°C. The flow rate was 0.6 mL/min. Phase A consisted of water and phase B was acetonitrile. Column separation for six ginsenosides (Rg1, Re, Rb1, Rc, Rb2, and Rd) was performed by gradient elution program: 0-12 min, 15-27% B; 12-15 min, 27-30% B; 15-16.5 min, 30-34% B; 16.5-21 min, 34-36% B; 21-24 min, 36-38% B; 24-28 min, 38-44% B; 28-29 min, 44-45% B; 29-29.5 min, 45-100% B; 29.5-32 min, 100% B; 32-32.5 min, 100-15% B; 32.5-35 min, 15% B **(Supplementary Fig. 1)**.

### UHPLC-MS/MS

Chromatographic analysis was performed on an Agilent 1290 infinity II. Chromatographic separation was performed on a ACQUITY UPLC BEH C18 column (2.1 × 10 mm, 1.7 μm). The column temperature was 35°C. The flow rate was 0.6 mL/min. Phase A consisted of water and 0.1% formic acid (v/v), and phase B was acetonitrile and 0.1% formic acid (v/v). Column separation for ginsenoside F11 was performed by gradient elution program: 0-14.5 min, 15-30% B; 14.5-15.5 min, 30-32% B; 15.5-18.5 min, 32-38% B; 18.5-24 min, 38-43% B; 24-27 min, 43-55% B; 27-31 min, 55% B; 31-33 min, 55-90% B; 33-38 min, 90% B; 38-38.1 min, 90-15% B; 38.1-43 min, 15% B (**Supplementary Fig. 2**). Mass spectrometric analysis was performed on a triple four-stage mass spectrometer AB Sciex QTRAP® 6500+ operated in negative ESI mode. The MS/MS parameters were optimized as follows: atomization temperature, 400°C; curtain gas, 30.0 psi; collision gas, 9.0 psi; ion spray voltage, -4.5 kV; ion source gas 1, 50.0 psi; ion source gas 2, 70.0 psi. Information about the MRM transition is shown in **Table 1**. Ginsenoside contents are shown in **Table 2**.

**Table 1.** MRM transition.

| **Analytes** | **Precursor ion** | **Production** | **Decluttering potential** | **Collision energy** |
| --- | --- | --- | --- | --- |
| Ginsenoside F11-1 | 845.26 | 799.40 | -65.00 | -32.00 |
| Ginsenoside F11-2 | 845.26 | 653.20 | -65.00 | -52.00 |
| Ginsenoside F11-3 | 845.263 | 491.20 | -65.00 | -60.00 |

**Table 2.** Information of ginsenoside contents of G1899.

| **Ginsenoside** | **Concentration in G1899 (mg/g)** |
| --- | --- |
| Rg1 | 1.30 |
| Re | 13.18 |
| F11 | 2.34 |
| Rb1 | 47.30 |
| Rc | 3.45 |
| Rb2 | 0.57 |
| Rd | 4.49 |

### G1899 administration

200 mg/kg of G1899 was orally administrated to 2-3-month-old mice daily for 4 weeks before performing the experiments as shown in Fig. 1.

### Acute stress

Lipopolysaccharide (LPS) is known to induce stress in several ways including oxidative stress, inflammatory responses, and cytokine expression (Asti et al., 2000; Couch et al., 2016; Zhao et al., 2023). Therefore, a single low-dose of 0.01 mg/kg LPS, a condition showing no abnormality in locomotor activity and overall animals’ health, was intraperitoneally injected to 2-3-month-old C57Bl6J female and male mice to induce acute stress. Saline injection was also used as a naïve control. 24 hours after LPS injection, we performed the experiments as described in as shown in **Fig. 1a**.

**Figure 1.**
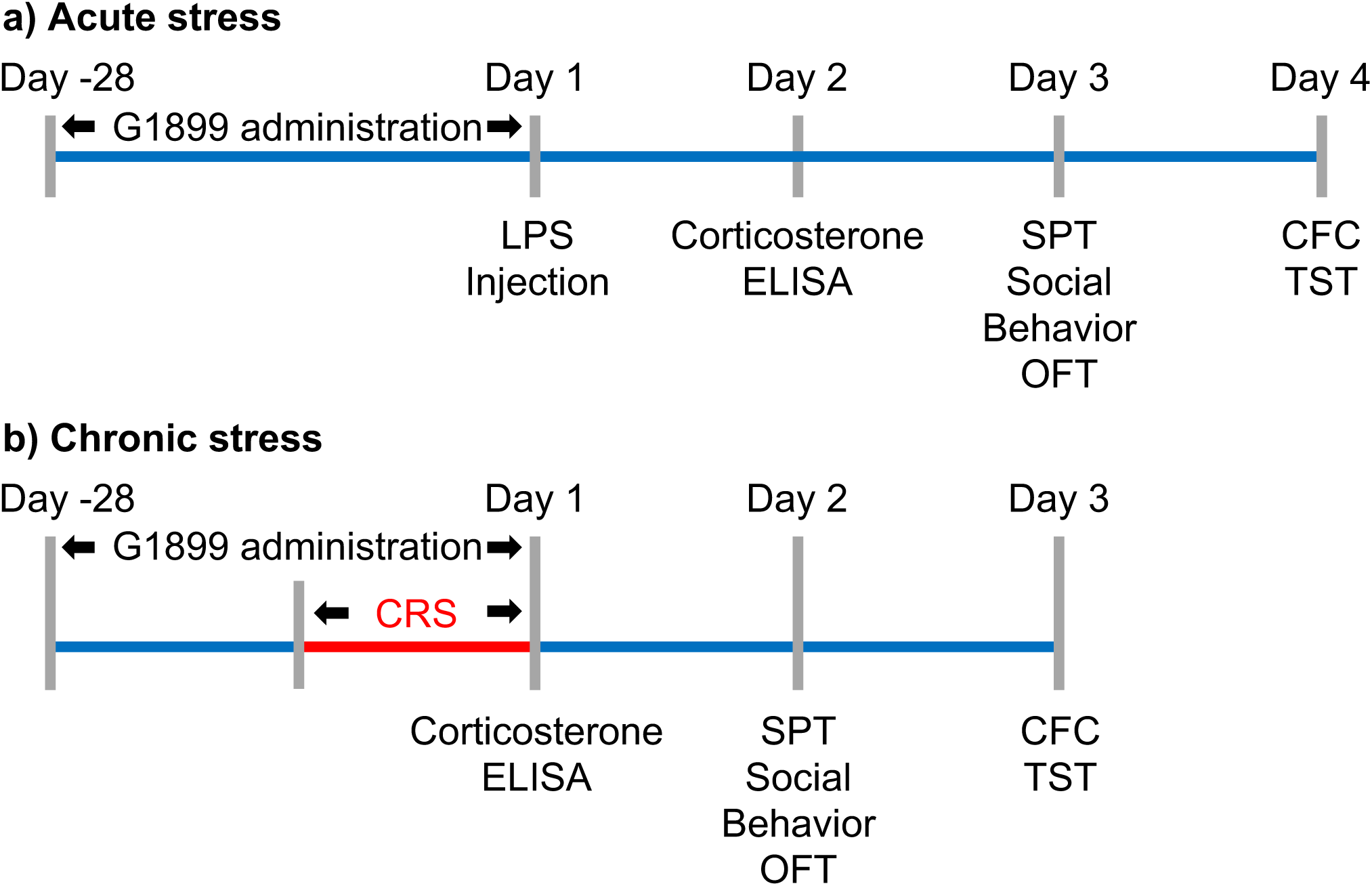
Experimental timeline of **a)** LPS-induced acute stress and **b)** chronic restraint stress (CRS) models. **a)** G1899 is daily administered to animals for 4 weeks. After G1899 administration, low-dose LPS (0.01mg/kg) is injected. Animals’ blood is collected to measure corticosterone levels using ELISA 24 hours after LPS injection. Multiple behavioral tests – sucrose preference test (SPT), reciprocal social interaction test, open-field test (OFT), contextual fear conditioning (CFC), and tail suspension test (TST) – are conducted on Day 3 and 4. **b)** G1899 is daily administered to animals for 4 weeks. 2 weeks after G1899 administration, CRS is applied to mice for 2 weeks. Blood collection and behavioral tests – SPT, reciprocal social interaction test, OFT, CFC, and TST – are carried out on Day 2 and 3.

### Chronic Restraint stress (CRS)

2-3-month-old CD1 (ICR) female and male mice were individually placed into 50 mL polypropylene conical tubes with multiple holes for ventilation, and they were exposed to restraint stress (2 hours/day) for 14 consecutive days as shown previously (Flowers et al., 2024). After restraint stress, mice were returned to the home cage. Following 2-week CRS, we conducted the experiments as shown in **Fig. 1b**.

### Corticosterone levels

Corticosterone levels in mice were measured as described previously (Flowers et al., 2024). 50 µl tail-tip whole blood samples were collected and centrifuged at 5,000 rpm for 10 minutes. 10 µl of serum was harvested and stored at −20°C until assay. 5 µl of serum samples in each condition was assayed using the DetectX® Corticosterone Multi-Format Enzyme-linked immunosorbent assay (ELISA) kit (Arbor Assays, Inc. K014-H).

### Sucrose preference test (SPT)

To measure motivation anhedonic behavior (a lack of pleasure), the SPT was conducted as shown previously (Kim et al., 2018b). Mice were be exposed to one bottle of water and one bottle of 1% sucrose. On day 1, the test mice were given the choice between water and a sucrose solution after initial habituation to two bottles of water for 24 hours. The following day (day 2), the total volume of liquid consumed from each bottle overnight was measured. The sucrose preference was calculated as the fraction of sucrose solution consumed compared to the total amount of solution consumed from both bottles. Bottles containing the sucrose solution was randomly placed on the left or right side of the cage. The lower sucrose preference indicates anhedonic behavior. Animals that did not consume water or sucrose were excluded from the analysis.

### Reciprocal social interaction

We performed the reciprocal social interaction test to examine social interaction between two freely moving mice. A test mouse (2-3-month-old) and a stranger mouse of the same genetic background that was previously housed in different cages were placed into the open field chamber (40 W x 40 L x 40 H cm) and allowed to explore freely for 20 minutes. Social behavior was recorded using a camera mounted overhead. For acute stress, social contacts were determined by the sniffing time that was defined as each instance in which a test mouse’s nose came within 2 cm toward a stranger mouse. The total number of reciprocal interactions and total duration of reciprocal interactions were measured manually and blindly by different investigators as described previously (Mendez-Vazquez et al., 2023). For chronic stress, social contacts were determined by the sniffing time that was defined as each instance in which a test mouse’s nose came within 0.25 cm toward a stranger mouse. Data were analyzed using the EthoVision XT tracking program (Noldus) to acquire the total number of reciprocal interactions and total duration of reciprocal interactions. Animals that escaped the open field chamber or showed aggressive behaviors (e.g. attacking a stranger mouse) were excluded from the analysis.

### Open field test (OFT)

We measured locomotor activity and anxiety-like behavior using the open field test as carried out previously (Flowers et al., 2024; Shou et al., 2018; Zaytseva et al., 2023). The test mouse was first placed in the center of the open field chamber (40 W x 40 L x 40 H cm) for 5 minutes. Animals were then allowed to explore the chamber for 20 minutes. The behavior was recorded by a video camera. A 20 x 20 cm center square was defined as the inside zone. Data were analyzed using the ANY-maze tracking program (Stoelting Co.) to acquire total traveled distance (locomotor activity) and time spent outside and inside (anxiety-like behavior). Animals that escaped the chamber or showed significantly decreased locomotor activity were excluded in analysis.

### Contextual fear conditioning (CFC)

Contextual fear conditioning (Habitest Modular System, Coulbourn Instrument) was carried out as described previously (Flowers et al., 2024; Kim et al., 2016; Lee et al., 2024; Shou et al., 2018). On Day 1, the test mouse was placed in a novel rectangular chamber with a grid floor. After a 3-minute baseline period, the test animal was given one shock (a 2-second, 0.5 mA shock) and stayed in the chamber for an additional one minute after the shock before being returned to the home cage for overnight. A contextual memory test was conducted on Day 2 in the same conditioning chamber for 3 minutes. Fear memory was determined by measuring the percentage of the freezing response (immobility excluding respiration and heartbeat) using an automated tracking program (FreezeFrame). Mice that showed no freezing were excluded from the analysis.

### Tail suspension test (TST)

We used the tail suspension test to examine depression-like behavior as described previously (Flowers et al., 2024; Kim et al., 2016; Kim et al., 2018b; Zaytseva et al., 2023). The test mouse was suspended by its tails from a rod suspended 20 cm above the tabletop surface with adhesive tape placed 1 cm from the tip of the tail. Animals were immobile when they exhibited no body movement and hung passively for > 3 seconds. The time during which mice remained immobile was quantified over a period of 6 minutes. The behavior was recorded by a video camera and, where data were analyzed using the ANY-maze tracking program to acquire immobility (depression-like behavior). Mice that successfully climbed their tails to escape were excluded from the analysis.

### Primary hippocampal neuron culture

Postnatal day 0 (P0) male and female CD-1 (ICR) pups were used to produce mouse hippocampal neuron cultures as shown previously (Sathler et al., 2022; Sathler et al., 2021; Sztukowski et al., 2018; Zaytseva et al., 2023). Hippocampi were isolated from P0 CD-1 (ICR) mouse brain tissues and digested with 10 U/mL papain (Worthington Biochemical Corp., LK003176). Mouse hippocampal neurons were plated on following poly lysine-coated glass bottom dishes (500,000 cells). Neurons were grown in Neurobasal Medium without phenol red (Thermo Fisher Scientific, 12348017) with B27 supplement (Thermo Fisher Scientific, 17504044), 0.5 mM Glutamax (Thermo Fisher Scientific, 35050061), and 1% penicillin/streptomycin (Thermo Fisher Scientific, 15070063).

### GCaMP Ca^2+^ Imaging

We measured spontaneous Ca^2+^ activity in cultured hippocampal neurons because It has been shown that networks of neurons in culture can produce spontaneous synchronized activity (Cohen et al., 2008). We infected 4 days *in vitro* (DIV) neurons with adeno-associated virus (AAV) expressing GCaMP8s under the neuron-specific human synapsin promoter (pGP-AAV-syn-jGCaMP8s-WPRE was a gift from GENIE Project (Addgene viral prep # 162374-AAV1; http://n2t.net/addgene:162374 ; RRID:Addgene_162374). We then measured Ca^2+^ activity in the soma of 14 DIV cultured hippocampal excitatory neurons with a modification of the previously described method (Kim et al., 2015a; Kim et al., 2015b; Kim and Ziff, 2014; Lee et al., 2024; Roberts et al., 2021; Sun et al., 2019; Sztukowski et al., 2018; Zaytseva et al., 2023). Glass-bottom dishes were mounted on a temperature-controlled stage on an Olympus IX73 microscope and maintained at 37°C and 5% CO_2_ using a Tokai-Hit heating stage and digital temperature and humidity controller. The images were captured right after 200 mg/L G1899 or saline was added to the media with a 10 ms exposure time and a total of 100 images were obtained with a one-second interval. F_min_ was determined as the minimum fluorescence value during the imaging. Total Ca^2+^ activity was obtained by 100 values of ΔF/F_min_ = (F_t_ – F_min_) / F_min_ in each image, and values of ΔF/F_min_ < 0.1 were rejected due to potential photobleaching. The average total Ca^2+^ activity in the control group was used to normalize total Ca^2+^ activity in each cell. The control group’s average total Ca^2+^ activity was compared to the experimental groups’ average as described previously (Kim et al., 2015a; Kim et al., 2015b; Kim and Ziff, 2014; Lee et al., 2024; Roberts et al., 2021; Sun et al., 2019; Sztukowski et al., 2018; Zaytseva et al., 2023).

### AAV infection in the mouse hippocampus and slice preparation

We virally expressed GCaMP using bilateral stereotaxic injection in the mouse hippocampus. Animals (3-month-old male and female C57Bl6 mice) were anesthetized with an intraperitoneal injection of a mixture of ketamine (100 mg/kg) and xylazine (10 mg/kg) and placed in a stereotactic frame. Anesthetic depth was confirmed with pedal response (foot retraction, response to non-damaging pressure of footpads using tweezers), ear twitch responses, and respiratory rates. Animal temperature was maintained with heating pads or warming gel packs. Once it was confirmed that the mice were properly anesthetized, the surgical field of the head of mice was aseptically prepared (shaved and prepped with betadine and alcohol). Animals were then placed in a stereotaxic frame (Stoelting). A small incision of the scalp was made with a sterile #10 surgical blade. With the aid of stereotaxic mounting equipment, a small hole was drilled in the bone using a high-speed drill and a dental bone drill bit, which has been sterilized. When the dura was exposed, a small pin hole was made, and a sterile syringe to inject 2 μl of AAV expressing GCaMP8f under the neuron-specific human synapsin promoter (pGP-AAV-syn-jGCaMP8f-WPRE was a gift from GENIE Project (Addgene viral prep # 162376-AAV1; http://n2t.net/addgene:162376 ; RRID:Addgene_162376) was lowered to the hippocampal CA1 area (Bregma coordinates: AP: − 1.95 mm, ML: ± 1.12 mm, DV: − 1.20 mm). During surgery, anesthetic depth was monitored every 5 minutes using pedal responses and respiration rates. After surgery, animals were allowed to recover from the anesthesia before being returned to their cages, and their health was closely monitored. Mice received analgesic doses of buprenorphine every 12 hours for one day after surgery. Buprenorphine was delivered by subcutaneous injection (0.1 mg/kg). Mice were monitored for any of the following signs of prolonged discomfort and pain: aggressiveness, hunched posture, failure to groom, awkward gait, vocalization, greater or less tissue coloration, eye discoloration, abnormal activity (usually less), hesitancy to move (especially in response to startle), water consumption, or food intake. We used transverse hippocampal slices (300 μm) containing the dorsal hippocampal CA1 area 2 weeks after the viral injection to ensure viral GCaMP8f expression. Hippocampal slices were cut and transferred to a holding chamber where slices will be maintained at 30°C and perfused with artificial cerebrospinal fluid (ACSF) continuously oxygenated with 95% O_2_ and 5% CO_2_ as shown previously (Flowers et al., 2024).

### GCaMP Ca^2+^ Imaging with glutamate uncaging

We carried out Ca^2+^ imaging with glutamate uncaging as shown previously (Zaytseva et al., 2023) in cultured hippocampal neurons and hippocampal slices right after 200 mg/L G1899 or saline was added to the media and ACSF, respectively. For glutamate uncaging in cultured neurons, 1 mM 4-methoxy-7-nitroindolinyl (MNI)-caged L-glutamate was added to the culture media, and epi-illumination photolysis (390 nm, 0.12 mW/mm^2^, 1 ms) was used for cultured neurons. For hippocampal slices, 0.25 mM MNI-caged L-glutamate was added to ACSF, and three stimulations of epi-illumination photolysis (390 nm, 0.12 mW/mm^2^, 1 ms, 50 ms interval) were applied. 1 μM tetrodotoxin (TTX) was added to prevent action potential-dependent network activity in culture media and ACSF. A baseline average of 20 frames (50 ms exposure) (F_0_) were captured prior to glutamate uncaging, and 50 more frames (50 ms exposure) were obtained after photostimulation. The fractional change in fluorescence intensity relative to baseline (ΔF/F_0_) was calculated. The average peak amplitude in the control group was used to normalize the peak amplitude in the G1899-treated group. The control group’s average peak amplitude was compared to the G1899-treated groups’ average.

### Statistical analysis

All behavior tests were blindly scored by more than two investigators. The Franklin A. Graybill Statistical Laboratory at Colorado State University was consulted for statistical analysis in the current study, including sample size determination, randomization, experiment conception and design, data analysis, and interpretation. We used the GraphPad Prism 10 software to determine statistical significance (set at p < 0.05). Grouped results of single comparisons were tested for normality with the Shapiro-Wilk normality or Kolmogorov-Smirnov test and analyzed using an unpaired two-tailed Student’s t-test when data are normally distributed. Differences between multiple groups were assessed by 2-way analysis of variance (ANOVA) with the Tukey test. The graphs were presented as mean ± Standard Deviation (SD). We discard data points that are located further than two SD above or below the average as an outlier.

## Results

### American ginseng (*Panax quinquefolius* L.) extract administration significantly decreases serum corticosterone levels only in chronically stressed animals

Stress activates the hypothalamic-pituitary-adrenal (HPA) axis, resulting in the release of glucocorticoid class stress hormones (Joels et al., 2018; McEwen, 2007; Phillips et al., 2006). Cortisol is the primary endogenous adrenal steroid in most mammals, including humans, whereas corticosterone is the primary adrenal corticosteroid in rodents (Raff et al., 2014; Raff et al., 1989; Usa et al., 2007; Yu et al., 2015). Stress can keep these hormone levels high, which in turn alters brain functions (Joels et al., 2018). We thus examined wither G1899 administration affected serum corticosterone levels in stressed animals. To induce stress in animals, we used two protocols, lipopolysaccharide (LPS) injection for acute stress and chronic restraint stress (CRS) for chronic stress. Both LPS injection and CRS are known to increase corticosterone levels in animals, which is associated with stress-induced behavioral changes (Marin et al., 2007; Russell et al., 2022). As shown previously (Russell et al., 2022), we found that LPS injection significantly increased serum corticosterone levels in mice when compared to saline-injected naïve mice (Naïve, 2394.10 ± 896.417 pg/ml and LPS, 4137.00 ± 690.41 pg/ml, p = 0.0014) (**Fig. 2a and Table 3**). Interestingly, G1899 administration significantly elevated serum corticosterone levels in LPS-injected animals (LPS + G1899, 5268.33 ± 1259.73 pg/ml, p = 0.045), whereas it had no effect on corticosterone levels in naïve mice (Naïve + G1899, 2375.82 ± 1106.97 pg/ml, p > 0.9999) (**Fig. 2a and Table 3**). Next, we found that CRS markedly elevated serum corticosterone levels in animals when compared to naïve mice (Naïve, 834.14 ± 637.27 pg/ml and CRS, 2460.42 ± 2546.52 pg/ml, p = 0.02) as reported previously (Flowers et al., 2024) (**Fig. 2b and Table 4**). Importantly, G1899 administration significantly reduced serum corticosterone levels in CRS animals (LPS + CRS, 877.28 ± 524.75 pg/ml, p = 0.0189) (**Fig. 2b and Table 4**). However, G1899 had no effect on corticosterone levels in naïve animals (Naïve + G1899, 680.02 ± 577.43 pg/ml, p = 9950) (**Fig. 2b and Table 4**). These findings show that both low-dose LPS injection and CRS are sufficient to increase stress hormone levels in animals, an indication of stress induction. However, G1899 administration significantly increases and decreases serum corticosterone levels in LPS-injected and CRS mice, respectively.

**Figure 2.**
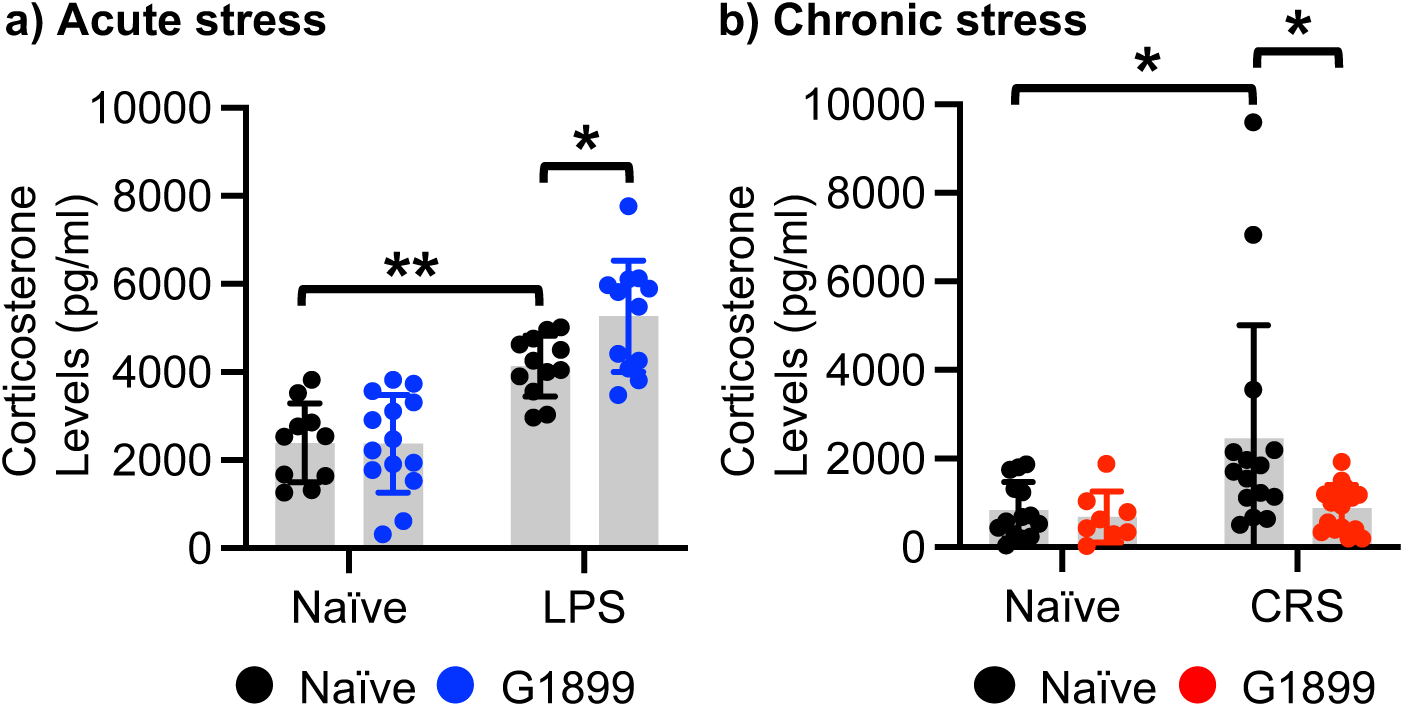
G1899 administration significantly decreases serum corticosterone levels only in chronic stressed animals. Summary of serum corticosterone levels in **a)** Naïve and LPS-induced acutely stressed animals in the presence or absence of G1899 (n = numbers of animals [numbers of females + numbers of males], Naïve = 10 [6 + 4], Naïve + G1899 = 14 [4 + 10], LPS = 12 [6 + 6], and LPS + G1899 = 12 [6 + 6]) and a) Naïve and CRS-induced chronically stressed animals in the presence or absence of G1899 (Naïve = 14 [9 + 5], Naïve + G1899 = 8 [5 + 3], CRS = 15 [9 + 6], and CRS + G1899 = 16 [10 + 6]). Two-way ANOVA. Tukey test, *p < 0.05 and **p < 0.01.

**Table 3.** Statistical analysis of serum corticosterone levels in Fig. 2a.

| LPS |  |  |  |  |  |
| --- | --- | --- | --- | --- | --- |
| ANOVA table | SS (Type III) | DF | MS | F (DFn, DFd) | P value |
| Interaction | 3908300 | 1 | 3908300 | F (1, 44) = 3.750 | P=0.0593 |
| Row Factor (Saline vs. G1899) | 63550485 | 1 | 63550485 | F (1, 44) = 60.97 | P<0.0001 |
| Column Factor (Naïve vs. LPS) | 3664984 | 1 | 3664984 | F (1, 44) = 3.516 | P=0.0674 |
| Residual | 45861574 | 44 | 1042308 |  |  |
| Tukey's multiple comparisons test | Predicted (LS) mean diff. |  | 95.00% CI of diff. |  | Adjusted P Value |
| Naïve:Saline vs. Naïve:G1899 | 18.18 |  | -1110 to 1147 |  | >0.9999 |
| Naïve:Saline vs. LPS:Saline | -1743 |  | -2910 to -575.7 |  | 0.0014 |
| Naïve:Saline vs. LPS:G1899 | -2874 |  | -4041 to -1707 |  | <0.0001 |
| Naïve:G1899 vs. LPS:Saline | -1761 |  | -2833 to -688.7 |  | 0.0004 |
| Naïve:G1899 vs. LPS:G1899 | -2892 |  | -3965 to -1820 |  | <0.0001 |
| LPS:Saline vs. LPS:G1899 | -1131 |  | -2244 to -18.49 |  | 0.0450 |

**Table 4.** Statistical analysis of serum corticosterone levels in Fig. 2b.

| CRS |  |  |  |  |  |
| --- | --- | --- | --- | --- | --- |
| ANOVA table | SS (Type III) | DF | MS | F (DFn, DFd) | P value |
| Interaction | 6271914 | 1 | 6271914 | F (1, 49) = 2.997 | P=0.0897 |
| Row Factor (Saline vs. G1899) | 10212953 | 1 | 10212953 | F (1, 49) = 4.881 | P=0.0319 |
| Column Factor (Naïve vs. CRS) | 9269346 | 1 | 9269346 | F (1, 49) = 4.430 | P=0.0405 |
| Residual | 102530333 | 49 | 2092456 |  |  |
| Tukey's multiple comparisons test |  |  |  |  |  |
|  | Predicted (LS) mean diff. | 95.00% CI of diff. |  | Adjusted P Value |  |
| Naïve:Saline vs. Naïve:G1899 | 154.1 | -1551 to 1859 |  | 0.9950 |  |
| Naïve:Saline vs. CRS:Saline | -1626 | -3056 to -196.7 |  | 0.0200 |  |
| Naïve:Saline vs. CRS:G1899 | -43.14 | -1451 to 1365 |  | 0.9998 |  |
| Naïve:G1899 vs. CRS:Saline | -1780 | -3465 to -96.20 |  | 0.0345 |  |
| Naïve:G1899 vs. CRS:G1899 | -197.3 | -1863 to 1469 |  | 0.9890 |  |
| CRS:Saline vs. CRS:G1899 | 1583 | 200.5 to 2966 |  | 0.0189 |  |

### American ginseng (*Panax quinquefolius* L.) extract administration induces anti-stress effects on acute stress-induced behavioral dysfunction in animals

LPS can cause a range of behavioral changes related to stress in humans and rodents, including anhedonia, depression-like behavior, and cognitive dysfunction (Cordeiro et al., 2019; van Heesch et al., 2013; Zhou et al., 2024). We thus carried out multiple behavioral assays to determine whether G1899 administration reversed acute stress-induced behavioral dysfunction in LPS-injected mice. First, we used the sucrose preference test (SPT) to determine anhedonia (reduced ability to experience pleasure). As shown previously (Arandelovic et al., 2024; Gu et al., 2018), we found a significant lower preference for sucrose following LPS injection, an indication of anhedonia, (Naïve, 0.70 ± 0.11 and LPS, 0.39 ± 0.17, p < 0.0001) (**Fig. 3a and Table 5**). However, G1899 significantly increased the preference for sucrose consumption in LPS-treated animals while it had no effect on the preference for sucrose consumption in naïve mice (Naïve + G1899, 0.75 ± 0.12, p = 08068, and LPS +G1899, 0.65 ± 0.16, p < 0.0001) (**Fig. 3a and Table 5**). This data suggests that G1899 is sufficient to reverse LPS-induced anhedonic depression-like behavior in animals.

**Figure 3.**
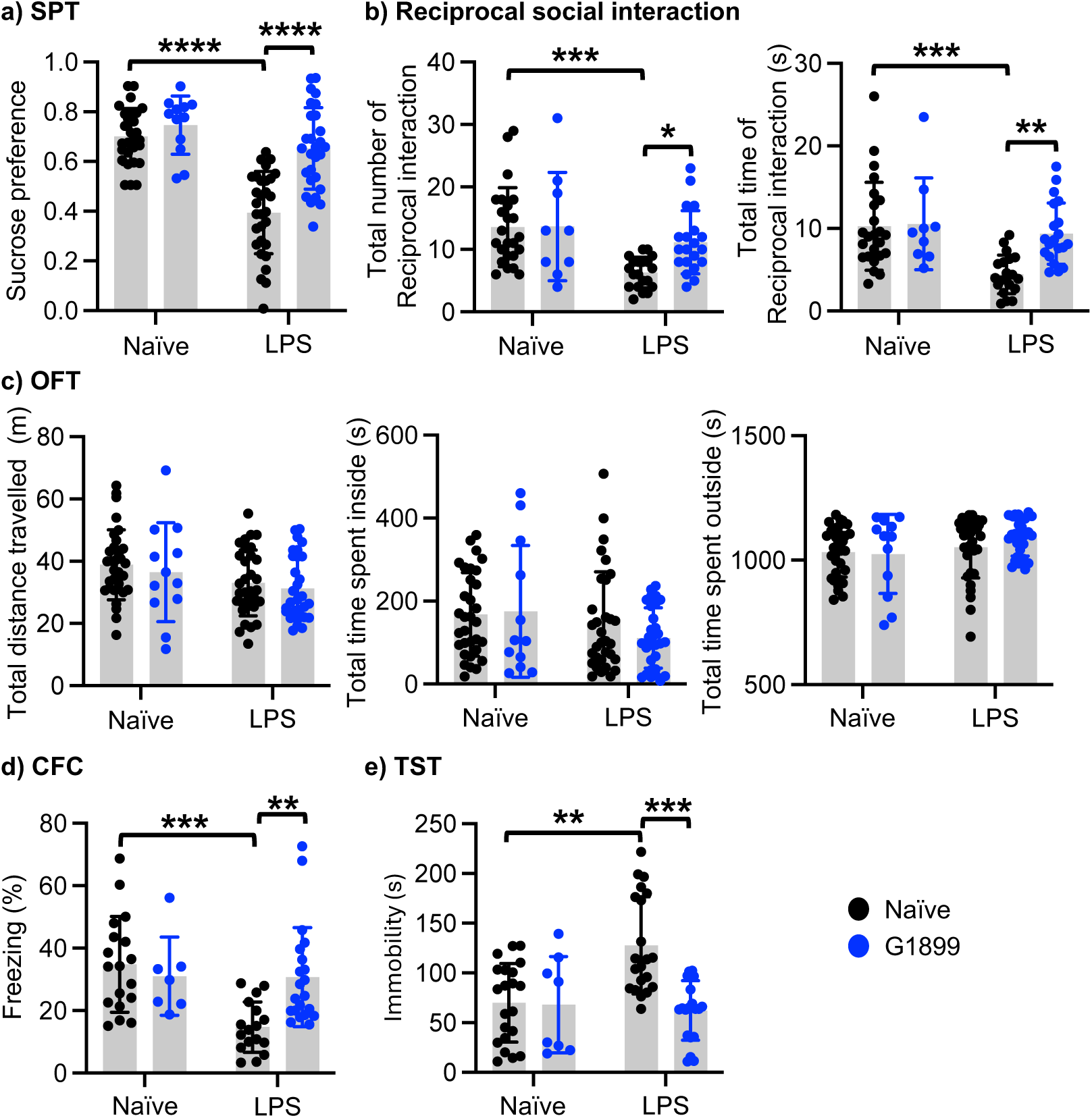
G1899 administration induces anti-stress effects on acute stress-induced behavioral dysfunction in animals. Summary of **a)** sucrose preference test (SPT) in each condition (n = numbers of animals [numbers of females + numbers of males], Naïve = 28 [16 + 12], Naïve + G1899 = 12 [5 + 7], LPS = 30 [14 + 16], and LPS + G1899 = 28 [12 + 16]), **b)** reciprocal social behavior test in each condition (Naïve = 24 [11 + 13], Naïve + G1899 = 9 [3 + 6], LPS = 20 [11 + 9], and LPS + G1899 = 20 [10 + 10]), **c)** open field test (OFT) in each condition (Naïve = 32 [16 + 16], Naïve + G1899 = 12 [5 + 7], LPS = 32 [16 + 16], and LPS + G1899 = 30 [13 + 17]), **d)** contextual fear conditioning (CFC) in each condition (Naïve = 18 [7 + 11], Naïve + G1899 = 7 [3 + 4], LPS = 17 [8 + 9], and LPS + G1899 = 21 [11 + 10]), and **e)** tail suspension test (TST) in each condition (Naïve = 21 [7 + 14], Naïve + G1899 = 8 [5 + 3], LPS = 21 [10 + 11], and LPS + G1899 = 20 [7 + 13]). Two-way ANOVA. Tukey test, *p < 0.05, **p < 0.01, ***p < 0.001, and ****p < 0.0001.

**Table 5.** Statistical analysis of the sucrose preference test in Fig. 3a.

| SPT |  |  |  |  |  |
| --- | --- | --- | --- | --- | --- |
| ANOVA table | SS (Type III) | DF | MS | F (DFn, DFd) | P value |
| Interaction | 0.2427 | 1 | 0.2427 | F (1, 94) = 11.32 | P=0.0011 |
| Row Factor (Saline vs. G1899) | 0.8467 | 1 | 0.8467 | F (1, 94) = 39.51 | P<0.0001 |
| Column Factor (Naïve vs. LPS) | 0.4920 | 1 | 0.4920 | F (1, 94) = 22.96 | P<0.0001 |
| Residual | 2.015 | 94 | 0.02143 |  |  |
| Tukey's multiple comparisons test |  |  |  |  |  |
|  | Predicted (LS) mean diff. | 95.00% CI of diff. |  | Adjusted P Value |  |
| Naïve:Saline vs. Naïve:G1899 | -0.04527 | -0.1774 to 0.08685 |  | 0.8068 |  |
| Naïve:Saline vs. LPS:Saline | 0.3064 | 0.2057 to 0.4070 |  | <0.0001 |  |
| Naïve:Saline vs. LPS:G1899 | 0.04744 | -0.05490 to 0.1498 |  | 0.6206 |  |
| Naïve:G1899 vs. LPS:Saline | 0.3516 | 0.2208 to 0.4824 |  | <0.0001 |  |
| Naïve:G1899 vs. LPS:G1899 | 0.09271 | -0.03941 to 0.2248 |  | 0.2634 |  |
| LPS:Saline vs. LPS:G1899 | -0.2589 | -0.3596 to -0.1583 |  | <0.0001 |  |

We conducted the reciprocal social interaction test to examine whether LPS injection induced social dysfunction and whether G1899 administration reversed LPS-induced social dysfunction. First, we found that LPS injection significantly decreased the total number of reciprocal interactions (Naïve, 13.58 ± 6.32 and LPS, 6.3 ± 2.45, p = 0.0003) and total time of reciprocal interactions (Naïve, 10.26 ± 5.34 seconds and LPS, 4.43 ± 2.33 seconds, p = 0.0002) in mice when compared to naïve controls, an indication of social dysfunction (**Fig. 3b and Table 6**). This suggests that LPS-induced acute stress significantly disrupted social behavior in animals. We next discovered that G1899 markedly increased the total number of reciprocal interactions (LPS + G1899, 11.15 ± 5.04, p = 0.0355) and total time of reciprocal interactions (LPS + G1899, 9.37 ± 3.71 seconds, p = 0.0028) in mice (**Fig. 3b and Table 6**). However, G1899 had no effect on social behavior in naïve animals (the total number of reciprocal interactions; Naïve + G1899, 13.67 ± 8.66, p > 0.9999 and total time of reciprocal interactions; Naïve + G1899, 10.56 ± 5.58 seconds, p = 0.9980) (**Fig. 3b and Table 6**). These findings suggest that G1899 can restore normal social behavior in acutely stressed animals.

**Table 6.** Statistical analysis of the reciprocal social interaction test in Fig. 3b.

|  |  |  |  |  |  |
| --- | --- | --- | --- | --- | --- |
| Total number of reciprocal interactions |  |  |  |  |  |
| ANOVA table | SS (Type III) | DF | MS | F (DFn, DFd) | P value |
| Interaction | 89.89 | 1 | 89.89 | F (1, 69) = 2.930 | P=0.0914 |
| Row Factor (Saline vs. G1899) | 379.9 | 1 | 379.9 | F (1, 69) = 12.39 | P=0.0008 |
| Column Factor (Naïve vs. LPS) | 96.28 | 1 | 96.28 | F (1, 69) = 3.139 | P=0.0809 |
| Residual | 2117 | 69 | 30.68 |  |  |
| Tukey's multiple comparisons test | Predicted (LS) mean diff. |  | 95.00% CI of diff. |  | Adjusted P Value |
| Naïve:Saline vs. Naïve:G1899 | -0.08333 |  | -5.783 to 5.616 |  | >0.9999 |
| Naïve:Saline vs. LPS:Saline | 7.283 |  | 2.869 to 11.70 |  | 0.0003 |
| Naïve:Saline vs. LPS:G1899 | 2.433 |  | -1.981 to 6.848 |  | 0.4724 |
| Naïve:G1899 vs. LPS:Saline | 7.367 |  | 1.514 to 13.22 |  | 0.0078 |
| Naïve:G1899 vs. LPS:G1899 | 2.517 |  | -3.336 to 8.370 |  | 0.6711 |
| LPS:Saline vs. LPS:G1899 | -4.850 |  | -9.461 to -0.2389 |  | 0.0355 |
| Total time of reciprocal interactions |  |  |  |  |  |
| ANOVA table | SS (Type III) | DF | MS | F (DFn, DFd) | P value |
| Interaction | 85.27 | 1 | 85.27 | F (1, 69) = 4.638 | P=0.0348 |
| Row Factor (Saline vs. G1899) | 195.2 | 1 | 195.2 | F (1, 69) = 10.61 | P=0.0017 |
| Column Factor (Naïve vs. LPS) | 108.5 | 1 | 108.5 | F (1, 69) = 5.901 | P=0.0177 |
| Residual | 1269 | 69 | 18.39 |  |  |
| Tukey's multiple comparisons test | Predicted (LS) mean diff. |  | 95.00% CI of diff. |  | Adjusted P Value |
| Naïve:Saline vs. Naïve:G1899 | -0.2972 |  | -4.710 to 4.115 |  | 0.9980 |
| Naïve:Saline vs. LPS:Saline | 5.833 |  | 2.415 to 9.251 |  | 0.0002 |
| Naïve:Saline vs. LPS:G1899 | 0.8933 |  | -2.525 to 4.311 |  | 0.9013 |
| Naïve:G1899 vs. LPS:Saline | 6.131 |  | 1.599 to 10.66 |  | 0.0037 |
| Naïve:G1899 vs. LPS:G1899 | 1.191 |  | -3.341 to 5.722 |  | 0.8999 |
| LPS:Saline vs. LPS:G1899 | -4.940 |  | -8.510 to -1.370 |  | 0.0028 |

As animals’ locomotor capacities can influence behavior, we carried out the open field test (OFT) to examine whether there were any abnormalities in animals’ locomotion in each condition. We found that there was no difference in total distance traveled, total time spent outside, and total time spent inside between the conditions (**Fig. 3c and Table 7**). Moreover, as time spent outside and inside can be interpreted as anxiety-like behavior (Zaytseva et al., 2023), normal behaviors in the OFT indicate that animals used in our study had normal locomotor activity and no anxiety-like behavior.

**Table 7.** Summary of the open field test in Fig. 3c.

|  | Naïve | LPS | Naïve + G1899 | LPS + G1899 |
| --- | --- | --- | --- | --- |
| Total distance traveled (m) | 38.87 ± 11.25 | 33.47 ± 10.54 | 36.47 ± 15.89 | 31.26 ± 10.29 |
| Total time spent outside (sec) | 1032.39 ± 100.74 | 1051.81 ± 122.63 | 1024.93 ± 158.51 | 1089.18 ± 72.77 |
| Total time spent inside (sec) | 167.61 ± 100.74 | 148.19 ± 122.63 | 175.07 ± 158.81 | 110.83 ± 72.77 |

It has been shown that low-dose LPS disrupts hippocampus-dependent fear memory (Pugh et al., 1998). We thus examined whether G1899 reversed LPS-induced hippocampus-dependent fear memory loss. To measure hippocampus-dependent fear memory, contextual fear conditioning (CFC) was used. We found that freezing was significantly lower in LPS-injected animals compared to naïve controls, an indication of fear memory loss (Naïve, 34.81 ± 15.31% and LPS, 14.74 ± 8.06%, p = 0.0003) (**Fig. 3d and Table 8**). We discovered that G1899 treatment significantly increased freezing in LPS-injected mice (LPS + G1899, 30.75 ± 15.86%, p = 0.0036) (**Fig. 3d and Table 8**). In contrast, G1899 administration in naïve animals was unable to alter fear memory (Naïve + G1899, 31.03 ± 12.54%, p = 0.9246) (**Fig. 3d and Table 8**). This data suggests that G1899 can restore normal hippocampus-dependent fear memory in acutely stressed animals.

**Table 8.**
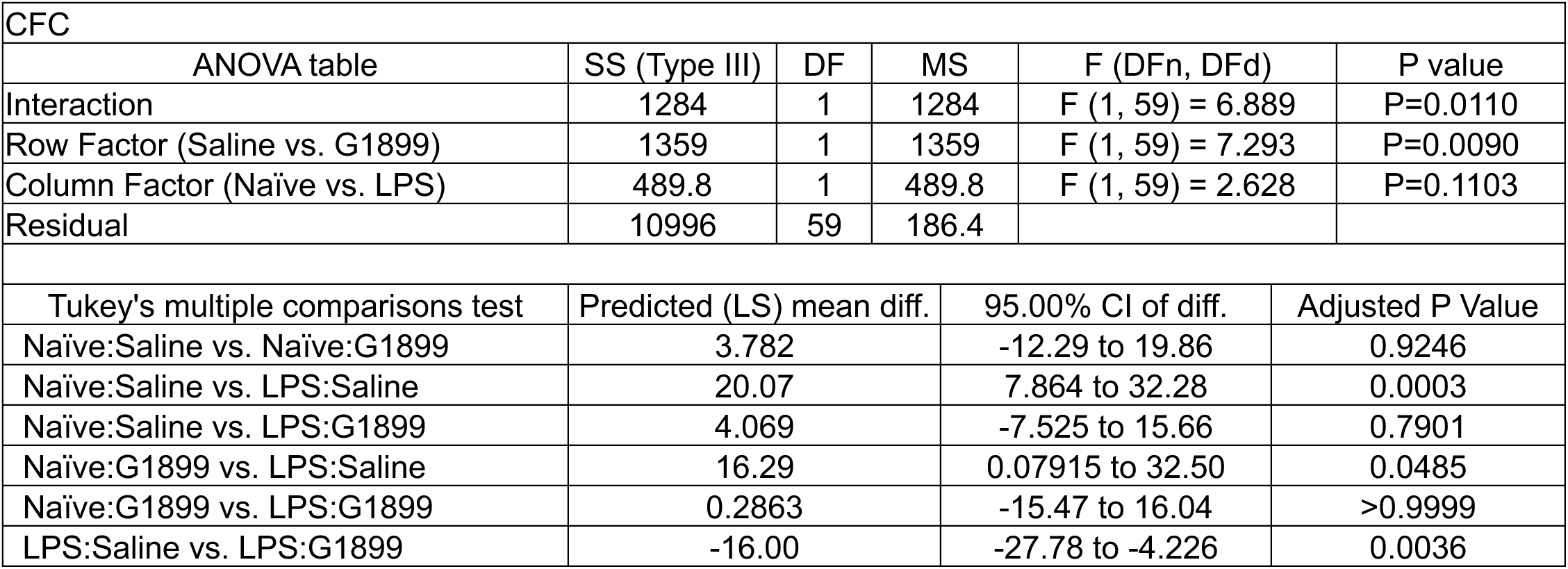
Statistical analysis of contextual fear conditioning in Fig. 3d.

| CFC |  |  |  |  |  |
| --- | --- | --- | --- | --- | --- |
| ANOVA table | SS (Type III) | DF | MS | F (DFn, DFd) | P value |
| Interaction | 1284 | 1 | 1284 | F (1, 59) = 6.889 | P=0.0110 |
| Row Factor (Saline vs. G1899) | 1359 | 1 | 1359 | F (1, 59) = 7.293 | P=0.0090 |
| Column Factor (Naïve vs. LPS) | 489.8 | 1 | 489.8 | F (1, 59) = 2.628 | P=0.1103 |
| Residual | 10996 | 59 | 186.4 |  |  |
| Tukey's multiple comparisons test | Predicted (LS) mean diff. |  | 95.00% CI of diff. |  | Adjusted P Value |
| Naïve:Saline vs. Naïve:G1899 | 3.782 |  | -12.29 to 19.86 |  | 0.9246 |
| Naïve:Saline vs. LPS:Saline | 20.07 |  | 7.864 to 32.28 |  | 0.0003 |
| Naïve:Saline vs. LPS:G1899 | 4.069 |  | -7.525 to 15.66 |  | 0.7901 |
| Naïve:G1899 vs. LPS:Saline | 16.29 |  | 0.07915 to 32.50 |  | 0.0485 |
| Naïve:G1899 vs. LPS:G1899 | 0.2863 |  | -15.47 to 16.04 |  | >0.9999 |
| LPS:Saline vs. LPS:G1899 | -16.00 |  | -27.78 to -4.226 |  | 0.0036 |

LPS injection is known to induce depression-like behavior in mice (Gu et al., 2018). We thus employed the tail suspension test (TST) to examine whether G1899 reversed LPS-induced depression-like behavior. As shown previously (Gu et al., 2018), LPS injection significantly elevated immobility in mice when compared to naïve controls, an indication of depression-like behavior (Naïve, 70.13 ± 39.55 seconds and LPS, 127.70 ± 48.85 seconds, p = 0.0002) (**Fig. 3e and Table 9**). Importantly, G1899 treatment markedly decreased immobility in LPS-injected animals, an indication of antidepressant effects (LPS + G1899, 62.35 ± 29.93 seconds, p < 0.0001) (**Fig. 3e and Table 9**). However, G1899 showed no effect on depression-like behavior in naïve mice (Naïve + G1899, 68.04 ± 48.55 seconds, p = 0.9993) (**Fig. 3e and Table 9**).

**Table 9.**
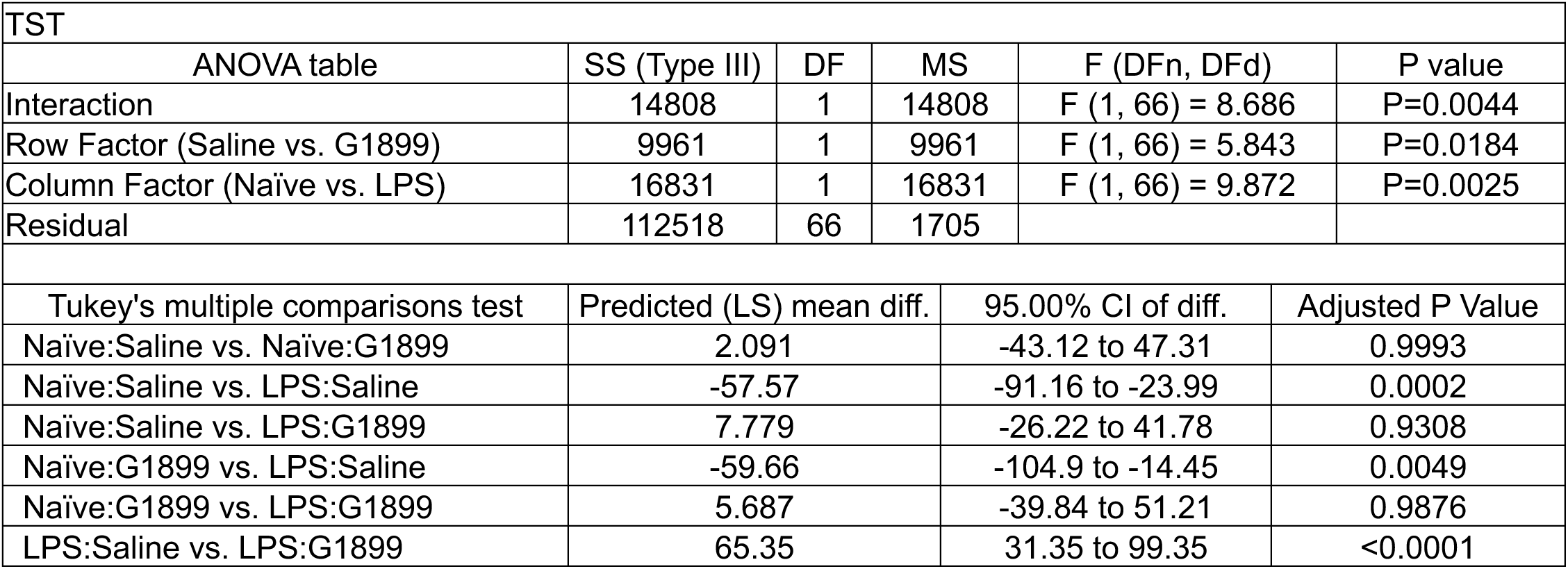
Statistical analysis of the tail suspension test in Fig. 3e.

| TST |  |  |  |  |  |
| --- | --- | --- | --- | --- | --- |
| ANOVA table | SS (Type III) | DF | MS | F (DFn, DFd) | P value |
| Interaction | 14808 | 1 | 14808 | F (1, 66) = 8.686 | P=0.0044 |
| Row Factor (Saline vs. G1899) | 9961 | 1 | 9961 | F (1, 66) = 5.843 | P=0.0184 |
| Column Factor (Naïve vs. LPS) | 16831 | 1 | 16831 | F (1, 66) = 9.872 | P=0.0025 |
| Residual | 112518 | 66 | 1705 |  |  |
| Tukey's multiple comparisons test |  |  |  |  |  |
|  | Predicted (LS) mean diff. | 95.00% CI of diff. |  | Adjusted P Value |  |
| Naïve:Saline vs. Naïve:G1899 | 2.091 | -43.12 to 47.31 |  | 0.9993 |  |
| Naïve:Saline vs. LPS:Saline | -57.57 | -91.16 to -23.99 |  | 0.0002 |  |
| Naïve:Saline vs. LPS:G1899 | 7.779 | -26.22 to 41.78 |  | 0.9308 |  |
| Naïve:G1899 vs. LPS:Saline | -59.66 | -104.9 to -14.45 |  | 0.0049 |  |
| Naïve:G1899 vs. LPS:G1899 | 5.687 | -39.84 to 51.21 |  | 0.9876 |  |
| LPS:Saline vs. LPS:G1899 | 65.35 | 31.35 to 99.35 |  | <0.0001 |  |

Our findings suggest that LPS-induced acute stress induces anhedonia, social dysfunction, fear memory loss, and depression-like behavior without altering locomotor activity, which are reversed by G1899 administration. Therefore, American ginseng (*Panax quinquefolius*) extract supplements likely have anti-stress effects in the LPS-induced acute stress model.

### American ginseng (*Panax quinquefolius* L.) extract administration induces anti-stress effects on chronic stress-induced behavioral dysfunction in animals

Chronic restraint stress (CRS) has been widely used as a model of chronic psychoemotional stress to induce depression- and anxiety-like behaviors, learning and memory deficits, social dysfunction, and hippocampal neuronal damage in mice (Huang et al., 2015; Ma et al., 2023; Park et al., 2018; Yun et al., 2010). We thus used CRS as a chronic stress model to test the G1899’s anti-stress effects. First, we used the SPT to determine anhedonia as CRS is shown to decrease sucrose preference in rodents (Rademacher and Hillard, 2007). As expected, we found a significant lower preference for sucrose following CRS, an indication of anhedonia, (Naïve, 0.71 ± 0.17 and CRS, 0.45 ± 0.11, p = 0.0002) (**Fig. 4a and Table 10**). Importantly, G1899 significantly increased the preference for sucrose consumption in CRS animals (CRS + G1899, 0.63 ± 0.17, p = 00402) (**Fig. 4a and Table 10**). We further found that G1899 had no effect on the preference for sucrose consumption in naïve mice (Naïve + G1899, 0.66 ± 0.13, p = 07553) (**Fig. 4a and Table 10**). These findings suggest that G1899 is sufficient to reverse CRS-induced anhedonic depression-like behavior in animals.

**Figure 4.**
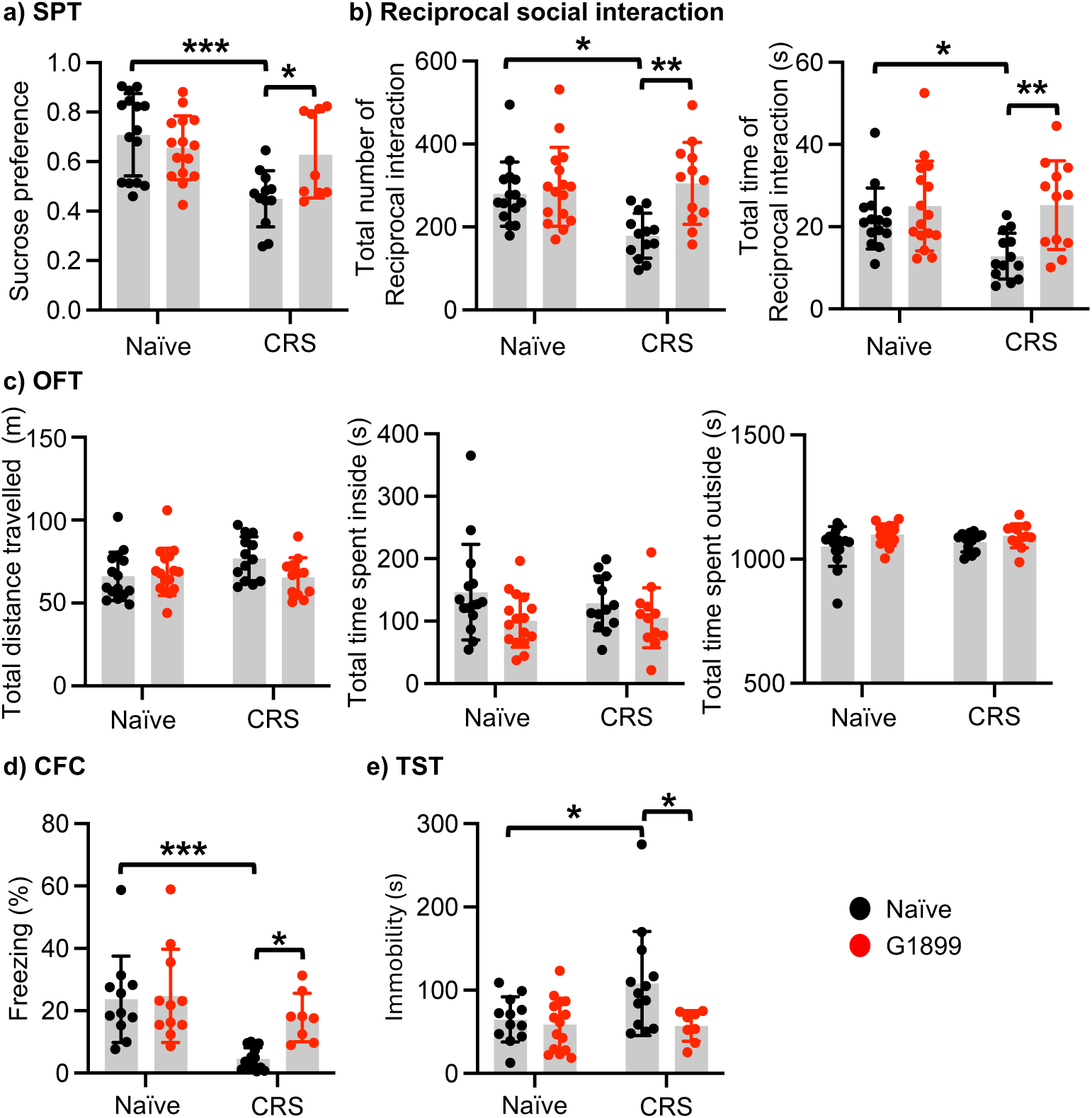
G1899 administration induces anti-stress effects on chronic stress-induced behavioral dysfunction in animals. Summary of **a)** sucrose preference test (SPT) in each condition (n = numbers of animals [numbers of females + numbers of males], Naïve = 15 [8 + 7], Naïve + G1899 = 15 [8 + 7], CRS = 12 [7 + 5], and CRS + G1899 = 9 [6 + 3]), **b)** reciprocal social behavior test in each condition (Naïve = 15 [10 + 5], Naïve + G1899 = 16 [9 + 7], CRS = 13 [7 + 6], and CRS + G1899 = 12 [8 + 4]), **c)** open field test (OFT) in each condition (Naïve = 15 [8 + 7], Naïve + G1899 = 16 [9 + 5], CRS = 13 [8 + 5], and CRS + G1899 = 12 [8 + 4]), **d)** contextual fear conditioning (CFC) in each condition (Naïve = 11 [5 + 6], Naïve + G1899 = 11 [7 + 4], CRS = 13 [8 + 5], and CRS + G1899 = 8 [5 + 3]), and **e)** tail suspension test (TST) in each condition (Naïve = 12 [7 + 5], Naïve + G1899 = 15 [10 + 5], CRS = 13 [8 + 5], and CRS + G1899 = 8 [5 + 3]). Two-way ANOVA. Tukey test, *p < 0.05, **p < 0.01, and ***p < 0.001.

**Table 10.** Statistical analysis of the sucrose preference test in Fig. 4a.

| SPT |  |  |  |  |  |
| --- | --- | --- | --- | --- | --- |
| ANOVA table | SS (Type III) | DF | MS | F (DFn, DFd) | P value |
| Interaction | 0.1621 | 1 | 0.1621 | F (1, 47) = 7.579 | P=0.0084 |
| Row Factor (Saline vs. G1899) | 0.2495 | 1 | 0.2495 | F (1, 47) = 11.67 | P=0.0013 |
| Column Factor (Naïve vs. CRS) | 0.04743 | 1 | 0.04743 | F (1, 47) = 2.217 | P=0.1431 |
| Residual | 1.005 | 47 | 0.02139 |  |  |
| Tukey's multiple comparisons test |  |  |  |  |  |
|  | Predicted (LS) mean diff. |  | 95.00% CI of diff. | Adjusted P Value |  |
| Naïve:Saline vs. Naïve:G1899 | 0.05291 |  | -0.08932 to 0.1951 | 0.7553 |  |
| Naïve:Saline vs. CRS:Saline | 0.2583 |  | 0.1074 to 0.4091 | 0.0002 |  |
| Naïve:Saline vs. CRS:G1899 | 0.08065 |  | -0.08358 to 0.2449 | 0.5626 |  |
| Naïve:G1899 vs. CRS:Saline | 0.2053 |  | 0.05448 to 0.3562 | 0.0038 |  |
| Naïve:G1899 vs. CRS:G1899 | 0.02774 |  | -0.1365 to 0.1920 | 0.9693 |  |
| CRS:Saline vs. CRS:G1899 | -0.1776 |  | -0.3494 to -0.005837 | 0.0402 |  |

Repeated stress can induce the development of social dysfunction (daSilva et al., 2021; Flowers et al., 2024). We thus conducted the reciprocal social interaction test to examine whether G1899 administration reversed CRS-induced social dysfunction. We revealed that CRS significantly decreased the total number of reciprocal interactions (Naïve, 279.53 ± 77.57 and CRS, 178.77 ± 54.33, p = 0.0128) and total time of reciprocal interactions (Naïve, 22.03 ± 7.43 seconds and CRS, 12.82 ± 5.59 seconds, p = 0.0452) in mice when compared to naïve controls, an indication of social dysfunction (**Fig. 4b and Table 11**), which is consistent with the previous findings (Flowers et al., 2024). Importantly, we further discovered that G1899 markedly increased the total number of reciprocal interactions (CRS + G1899, 305.08 ± 98.74, p = 0.0023) and total time of reciprocal interactions (CRS + G1899, 25.24 ± 10.80 seconds, p = 0.0061) in mice (**Fig. 4b and Table 11**). In contrast, G1899 had no effect on social behavior in naïve animals (the total number of reciprocal interactions; Naïve + G1899, 296.94 ± 95.30, p = 0.9378 and total time of reciprocal interactions; Naïve + G1899, 25.05 ± 10.95 seconds, p = 0.7869) (**Fig. 4b and Table 11**). This data suggests that G1899 can restore normal social behavior in chronically stressed animals.

**Table 11.** Statistical analysis of the reciprocal social interaction test in Fig. 4b.

|  |  |  |  |  |  |
| --- | --- | --- | --- | --- | --- |
| Total number of reciprocal interactions |  |  |  |  |  |
| ANOVA table | SS (Type III) | DF | MS | F (DFn, DFd) | P value |
| Interaction | 40983 | 1 | 40983 | F (1, 52) = 5.868 | P=0.0189 |
| Row Factor (Saline vs. G1899) | 29639 | 1 | 29639 | F (1, 52) = 4.244 | P=0.0444 |
| Column Factor (Naïve vs. CRS) | 71366 | 1 | 71366 | F (1, 52) = 10.22 | P=0.0024 |
| Residual | 363144 | 52 | 6984 |  |  |
| Tukey's multiple comparisons test |  |  |  |  |  |
|  | Predicted (LS) mean diff. |  |  | 95.00% CI of diff. | Adjusted P Value |
| Naïve:Saline vs. Naïve:G1899 | -17.40 |  |  | -97.12 to 62.31 | 0.9378 |
| Naïve:Saline vs. CRS:Saline | 100.8 |  |  | 16.72 to 184.8 | 0.0128 |
| Naïve:Saline vs. CRS:G1899 | -25.55 |  |  | -111.5 to 60.35 | 0.8589 |
| Naïve:G1899 vs. CRS:Saline | 118.2 |  |  | 35.35 to 201.0 | 0.0022 |
| Naïve:G1899 vs. CRS:G1899 | -8.146 |  |  | -92.85 to 76.55 | 0.9941 |
| CRS:Saline vs. CRS:G1899 | -126.3 |  |  | -215.1 to -37.52 | 0.0023 |
| Total time of reciprocal interactions |  |  |  |  |  |
| ANOVA table | SS (Type III) | DF | MS | F (DFn, DFd) | P value |
| Interaction | 305.4 | 1 | 305.4 | F (1, 52) = 3.757 | P=0.0580 |
| Row Factor (Saline vs. G1899) | 281.1 | 1 | 281.1 | F (1, 52) = 3.458 | P=0.0686 |
| Column Factor (Naïve vs. CRS) | 825.2 | 1 | 825.2 | F (1, 52) = 10.15 | P=0.0024 |
| Residual | 4228 | 52 | 81.30 |  |  |
| Tukey's multiple comparisons test |  |  |  |  |  |
|  | Predicted (LS) mean diff. |  |  | 95.00% CI of diff. | Adjusted P Value |
| Naïve:Saline vs. Naïve:G1899 | -3.026 |  |  | -11.63 to 5.575 | 0.7869 |
| Naïve:Saline vs. CRS:Saline | 9.211 |  |  | 0.1430 to 18.28 | 0.0452 |
| Naïve:Saline vs. CRS:G1899 | -3.217 |  |  | -12.49 to 6.052 | 0.7937 |
| Naïve:G1899 vs. CRS:Saline | 12.24 |  |  | 3.301 to 21.17 | 0.0035 |
| Naïve:G1899 vs. CRS:G1899 | -0.1908 |  |  | -9.330 to 8.948 | >0.9999 |
| CRS:Saline vs. CRS:G1899 | -12.43 |  |  | -22.01 to -2.848 | 0.0061 |

Next, we carried out OFT to examine animals’ locomotion in each condition. We discovered no difference in total distance traveled, total time spent outside, and total time spent inside between the conditions, an indication of normal locomotor activity and no anxiety-like behavior (**Fig. 4c and Table 12**).

**Table 12.** Summary of the open field test in Fig. 4c.

|  | Naïve | CRS | Naïve + G1899 | CRS + G1899 |
| --- | --- | --- | --- | --- |
| Total distance traveled (m) | 66.12 ± 14.56 | 76.80 ± 13.21 | 68.80 ± 14.20 | 65.49 ± 11.77 |
| Total time spent outside (sec) | 1051.16 ± 79.85 | 1067.16 ± 37.95 | 1098.34 ± 43.45 | 1093.54 ± 48.52 |
| Total time spent inside (sec) | 146.49 ± 76.65 | 128.31 ± 43.53 | 100.72 ± 42.54 | 105.42 ± 47.90 |

Repeated stress can impair learning and memory (Kim and Kim, 2023). In fact, it has been shown that CRS disrupts hippocampus-dependent fear memory in mice (Flowers et al., 2024; Yun et al., 2010). We thus examined whether G1899 reversed CRS-induced hippocampus-dependent fear memory loss using CFC. We revealed that freezing was significantly lower in CRS mice compared to naïve controls, an indication of fear memory loss (Naïve, 23.65 ± 13.87% and CRS, 4.56 ± 3.54%, p = 0.0008) (**Fig. 4d and Table 13**). We discovered that G1899 treatment significantly improved fear memory in CRS mice (CRS + G1899, 17.86 ± 7.79%, p = 0.0495) (**Fig. 4d and Table 13**). In contrast, G1899 administration in naïve animals had no effect on fear learning and memory (Naïve + G1899, 24.76 ± 14.94%, p = 0.9952) (**Fig. 4d and Table 13**). This data suggests that G1899 can restore normal hippocampus-dependent fear memory in CRS mice.

**Table 13.** Statistical analysis of contextual fear conditioning in Fig. 4d.

| CFC |  |  |  |  |  |
| --- | --- | --- | --- | --- | --- |
| ANOVA table | SS (Type III) | DF | MS | F (DFn, DFd) | P value |
| Interaction | 387.1 | 1 | 387.1 | F (1, 39) = 3.191 | P=0.0818 |
| Row Factor (Saline vs. G1899) | 1760 | 1 | 1760 | F (1, 39) = 14.51 | P=0.0005 |
| Column Factor (Naïve vs. CRS) | 541.6 | 1 | 541.6 | F (1, 39) = 4.465 | P=0.0411 |
| Residual | 4731 | 39 | 121.3 |  |  |
| Tukey's multiple comparisons test |  |  |  |  |  |
|  | Predicted (LS) mean diff. |  | 95.00% CI of diff. | Adjusted P Value |  |
| Naïve:Saline vs. Naïve:G1899 | -1.114 |  | -13.72 to 11.49 | 0.9952 |  |
| Naïve:Saline vs. CRS:Saline | 19.09 |  | 6.980 to 31.20 | 0.0008 |  |
| Naïve:Saline vs. CRS:G1899 | 5.786 |  | -7.948 to 19.52 | 0.6732 |  |
| Naïve:G1899 vs. CRS:Saline | 20.20 |  | 8.094 to 32.31 | 0.0004 |  |
| Naïve:G1899 vs. CRS:G1899 | 6.900 |  | -6.833 to 20.63 | 0.5387 |  |
| CRS:Saline vs. CRS:G1899 | -13.30 |  | -26.58 to -0.02141 | 0.0495 |  |

Repeated stress is a known trigger for depression in humans and depression-like behavior in animals (Richter-Levin and Xu, 2018; Tafet and Nemeroff, 2016). Indeed, CRS is shown to induce depression-like behavior in rodents (Flowers et al., 2024; Mao et al., 2022). We thus employed TST to examine whether G1899 reversed CRS-induced depression-like behavior. We found that CRS significantly elevated immobility in animals when compared to naïve controls, an indication of depression-like behavior (Naïve, 64.78 ± 27.13 seconds and LPS, 108.09 ± 62.54 seconds, p = 0.0483) (**Fig. 4e and Table 14**). Importantly, G1899 treatment markedly decreased immobility in CRS animals, an indication of antidepressant effects (CRS + G1899, 57.04 ± 18.41 seconds, p = 0.0349) (**Fig. 4e and Table 14**). In contrast, G1899 showed no effect on depression-like behavior in naïve animals (Naïve + G1899, 58.69 ± 31.66 seconds, p = 0.9796) (**Fig. 4e and Table 14**).

**Table 14.** Statistical analysis of the tail suspension test in Fig. 4e.

| TST |  |  |  |  |  |
| --- | --- | --- | --- | --- | --- |
| ANOVA table | SS (Type III) | DF | MS | F (DFn, DFd) | P value |
| Interaction | 5745 | 1 | 5745 | F (1, 44) = 3.539 | P=0.0666 |
| Row Factor (Saline vs. G1899) | 4930 | 1 | 4930 | F (1, 44) = 3.037 | P=0.0884 |
| Column Factor (Naïve vs. CRS) | 9279 | 1 | 9279 | F (1, 44) = 5.716 | P=0.0212 |
| Residual | 71432 | 44 | 1623 |  |  |
| Tukey's multiple comparisons test | Predicted (LS) mean diff. |  | 95.00% CI of diff. |  | Adjusted P Value |
| Naïve:Saline vs. Naïve:G1899 | 6.090 |  | -35.58 to 47.76 |  | 0.9796 |
| Naïve:Saline vs. CRS:Saline | -43.31 |  | -86.38 to -0.2423 |  | 0.0483 |
| Naïve:Saline vs. CRS:G1899 | 7.746 |  | -41.36 to 56.85 |  | 0.9746 |
| Naïve:G1899 vs. CRS:Saline | -49.40 |  | -90.16 to -8.633 |  | 0.0119 |
| Naïve:G1899 vs. CRS:G1899 | 1.656 |  | -45.44 to 48.75 |  | 0.9997 |
| CRS:Saline vs. CRS:G1899 | 51.05 |  | 2.713 to 99.40 |  | 0.0349 |

These findings suggest that repeated stress induces anhedonia, social dysfunction, fear memory loss, and depression-like behavior without altering locomotor activity, which are reversed by G1899 administration. Therefore, American ginseng (*Panax quinquefolius*) extract supplements likely have anti-stress effects in the chronic stress mouse model

### American ginseng (*Panax quinquefolius* L.) extract significantly reduces the activity in hippocampal neurons by suppressing synaptic activity

Ginseng ginsenosides are shown to decrease glutamatergic NMDARs (Kim et al., 2002; Kim et al., 2004), which is thought to be critical for ginseng’s beneficial effects. Moreover, NMDAR antagonists have been used to treat several diseases including depression and Alzheimer’s disease (Beaurain et al., 2024). As NMDAR inhibition reduces neuronal activity by suppressing excitatory synaptic activity, we first examined whether G1899 affected glutamatergic activity in cultured hippocampal neurons using Ca^2+^ imaging with glutamate uncaging. For Ca^2+^ imaging, a genetically encoded Ca^2+^ indicator, GCaMP, was used to measure somatic Ca^2+^ activity following glutamate release in cultured hippocampal excitatory neurons in the presence or absence of G1899. Given that neuronal Ca^2+^ is the secondary messenger responsible for transmitting depolarization status and synaptic activity (Gleichmann and Mattson, 2011), we carried out somatic Ca^2+^ imaging with glutamate uncaging in cultured mouse hippocampal neurons to measure glutamatergic activity.

We treated 14 DIV hippocampal cultures with G1899 and measured glutamate-induced Ca^2+^ signals. Glutamatergic activity was significantly lower in G1899-treated neurons than control cells (CTRL) (CTRL, 1.000 ± 0.374 ΔF/F_0_ and G1899, 0.802 ± 0.327 ΔF/F_0_, p = 0.0005. t = 3.548, df = 157) **(Fig. 5a**). We also measured glutamatergic activity of hippocampal cells in brain slices that have intact neural circuits. As shown in cultured neurons, G1899 treatment significantly reduced glutamate-induced Ca^2+^ signals in neurons from hippocampal slices when compared to control cells (CTRL) (CTRL, 1.000 ± 0.366 ΔF/F_0_ and G1899, 0.342 ± 0.298 ΔF/F_0_, p < 0.0001, t = 9.462, df = 90) (**Fig. 5b**). Finally, we determined whether G1899 decreased neuronal activity in cultured hippocampal neurons. As somatic Ca^2+^ levels in neurons indicate neuronal activity (Gleichmann and Mattson, 2011), we measured spontaneous somatic Ca^2+^ activity using GCaMP without TTX in 12-14 DIV cultured hippocampal excitatory neurons in the presence or absence of G1899. We measured spontaneous Ca^2+^ activity right after G1899 was treated and found a significant reduction in Ca^2+^ activity in G1899-treated neurons compared to control cells (CTRL) (CTRL, 1.000 ± 0.538 ΔF/F_min_ and G1899, 0.571 ± 0.505 ΔF/F_min_, p < 0.0001, t = 5.467, df = 191) (**Fig. 5c**). This data demonstrates that G1899 treatment significantly reduces neuronal Ca^2+^ activity in cultured hippocampal cells. These findings suggest that G1899 treatment significantly reduces the activity in hippocampal neurons by suppressing glutamatergic activity.

**Figure 5.**
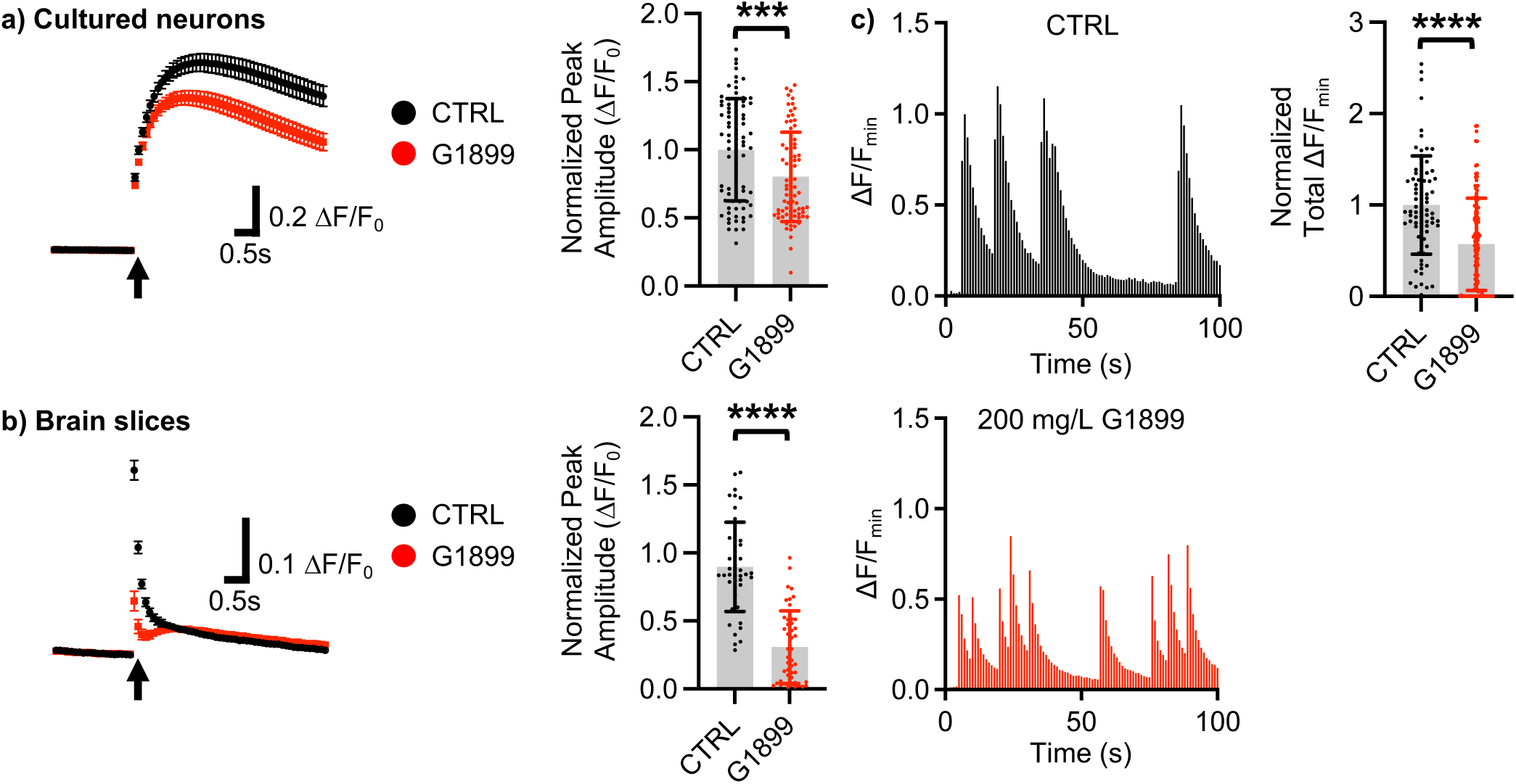
G1899 significantly reduces the activity in hippocampal neurons by suppressing synaptic activity. Average traces of virally expressed GCaMP8s signals and summary data of normalized peak amplitude in saline-(CTRL) and G1899-treated cells **a)** in cultures (n = number of neurons from 2 independent cultures, CTRL = 80 and G1899 = 79, ***p < 0.001, Two-tailed Student’s t-test) and **b)** hippocampal slices (n = number of neurons from 2 independent slices, CTRL = 46 and G1899 = 46, ****p < 0.0001, Two-tailed Student’s t-test). An arrow indicates photostimulation. **c)** Representative traces of GCaMP8s signals in cultured hippocampal cells and summary data of normalized total Ca^2+^ activity in saline-(CTRL) and G1899-treated cells (n = number of neurons from 2 independent cultures, CTRL = 78 and G1899 = 115, ****p < 0.0001, Two-tailed Student’s t-test).

## Discussion

Multiple studies have shown that ginseng likely protects the brain and enhances its function in pathological conditions. There is a compelling need for a more comprehensive understanding of ginseng’s role in the physiological condition because many individuals without specific diseases seek to improve their health by incorporating ginseng into their routines. Among many pathological conditions reversed by ginseng, it is suggested to increase long-term resistance to stress (Lee et al., 2021a; Xu et al., 2017). Chronic stress disrupts brain functions, contributing to the development of anxiety, depression, cognitive decline, and social dysfunction (daSilva et al., 2021; Kessler, 1997; Lupien et al., 2009). Nonetheless, previous research of ginseng’s effects on stress has been mainly focused on depression-like behavior in animals. Therefore, it is urgently needed to investigate whether ginseng can reverse a wide range of stress-induced pathological behaviors in animals. Here, we use both acute and chronic stress model mice and perform multiple behavioral assays to comprehensively examine whether ginseng supplements can induce the anti-stress effects. Indeed, we have found that daily administration of 200 mg/kg American ginseng for 4 weeks in mice significantly reverses acute and chronic stress-induced behavioral dysfunctions including anhedonia, social dysfunction, memory loss, and depression-like behavior. Moreover, our data suggests that American ginseng treatment reduces neuronal activity by suppressing glutamatergic activity in hippocampal neurons, the same effects on medications used for many neurological disorders (e.g. ketamine and memantine). Our findings thus likely provide new insight into the treatment of ginseng on stress-associated neurological diseases.

The roots of *Panax quinquefolius* L. (American ginseng) contains ginsenosides such as Rg1, Re, F11, Rb1, RC, Rb2, and Rd (**Table 2**) (Qi et al., 2011). Previous studies have investigated the effects of ginseng components on brain function, and several studies have shown their permeability across the blood-brain barrier (BBB) and the underlying mechanisms. For instance, ginsenoside Rb1 can pass the BBB via glucose transporter 1 (Wang et al., 2018). Moreover, the brain uptake of ginsenosides Rg1, Re, Rd and Rb1 has been found to be mediated by A1 adenosine receptors (A1Rs) (Liang et al., 2020). Additionally, following intravenous administration of fluorescent-labeled gintonin, a ginseng-derived G-protein-coupled lysophosphatidic acid (LPA) receptor ligand, gintonin is found in endothelial cells, neurons and glial cells in the brain (Kim et al., 2018a). Therefore, ginseng extract supplements can directly affect brain functions. Ginseng can also affect brain functions by influencing various cells within the central nervous system, including neurons, oligodendrocytes, astrocytes, and microglia (Shin et al., 2024). Moreover, it can affect neurotransmitter levels, synaptic receptors, and synaptic proteins at the molecular levels (Shin et al., 2024). Due to these complex cellular and molecular mechanisms of ginseng, it is difficult to understand how ginseng influences brain functions. One potential important mechanism is likely to be ginseng ginsenoside-induced NMDAR antagonism (Kim et al., 2002; Kim et al., 2004). In fact, NMDAR blockers have been widely used in clinics to treat various brain disorders, including stroke, epilepsy, Alzheimer’s disease, Huntington’s disease, schizophrenia, depression, and neuropathic pain (Beaurain et al., 2024). Importantly, several NMDAR antagonists such as dextromethorphan, esketamine (the S enantiomer form of ketamine), phencyclidine (PCP, also known as angel dust by drug addicts), thienylcyclohexylpiperidine (TCP), dizocilpine (MK-801), memantine, and fluoroethylnormemantine (FENM) have been approved by the Food and Drug Administration (FDA). Therefore, ginseng ginsenosides may work as NMDAR antagonists to induce the anti-stress effects. Most importantly, we have shown that low-dose ketamine treatment reverses CRS-induced social dysfunction, hippocampus-dependent fear memory loss, and depression-like behavior in both female and male mice (Flowers et al., 2024), which phenocopies the G1899 effects on mice. Importantly, ginsenoside Rg3 and its derivative Rk1 are known to inhibit NMDARs (Kim et al., 2002; Kim et al., 2004; Ryoo et al., 2020) although it is not abundant in American ginseng. However, Rb1 and Rg1, more abundant in American ginseng compared to Korea red ginseng, are known to reduce glutamatergic activity in hippocampal neurons (Lee et al., 2000). Rb1 and Rg1 are known to be BBB-permeable (Liang et al., 2020; Wang et al., 2018), whereas the BBB permeability of Rg3 has not been fully established, but it prevents BBB disruption in neurological disorders (Park et al., 2014). Therefore, it is possible that Rb1 and Rg1 from American ginseng can be new NMDAR antagonists to induce the anti-stress effects in mice.

Ginseng can help regulate the HPA axis, which is the body’s main stress response system (Lee and Rhee, 2017). When subjected to stress, the stress hormone, cortisol, is secreted to counteract stress and maintain homeostasis. However, prolonged cortisol secretion results in immunosuppression. Cortisol is produced and regulated by the HPA axis (Lee and Rhee, 2017). It has been shown that Rg1 alleviates anhedonia, hopelessness and improved sleep disruption through the modulation levels of corticosterone, testosterone, androgen receptor (AR), and glucocorticoid receptor (GR) in the chronic-unpredictable-mild-stress model and the gonadectomized model (Mou et al., 2017). Ginseng extracts also induce an antidepressant-like effect by inhibiting the HPA axis in depressed mice (Choi et al., 2018). Therefore, the G1899-induced reduction of serum corticosterone levels in CRS mice is likely mediated by suppressing the HPA axis (**Fig. 2b**). However, we have found that G1899 significantly elevates corticosterone levels in LPS-injected mice (**Fig. 2a**). LPS triggers a strong immune response by activating the NFκB pathway, resulting in the release of proinflammatory cytokines and stimulates an innate immune response (Park and Lee, 2013). In general, ginseng inhibits the NF-κB pathway, leading to anti-inflammatory effects (Kim et al., 2017). However, excessive consumption of ginseng may cause overactivation of the NF-κB pathway, which can lead to an overactive immune system and inflammatory responses (Deng et al., 2023). It is thus possible that 200 mg/kg G1899 administration is excessive in LPS-injected mice, which may lead to overactivation of inflammatory pathway, resulting in elevation of corticosterone levels. Importantly, these results provide a theoretical basis for improving ginseng quality and guidelines for the scientific consumption of ginseng. However, the specific component that may prevent side effects of ginseng overuse in the conditions related to LPS-induced stress still needs further study.

In summary, we reveal that both acute and chronic stress induce behavioral dysfunction, including depression-like behavior, anhedonia, social dysfunction, and fear memory loss in animals. We further demonstrate that G1899 treatment significantly reverses stress-induced behavioral abnormalities in animals. Our data also suggests that G1899 decreases neuronal activity in hippocampal neurons by reducing glutamatergic activity. These findings thus suggest that G1899 supplement can be protective against both acute and chronic stress in mice by suppressing neuronal and synaptic activity.

## Acknowledgments

We thank members of the Kim laboratory for their generous support. This work is supported by Student Experiential Learning Grants and College Research Council Shared Research Program from College of Veterinary Medicine and Biomedical Sciences, Colorado State University and a research grant from Korea Ginseng Corporation.

---

**Supplementary Figure 1.**
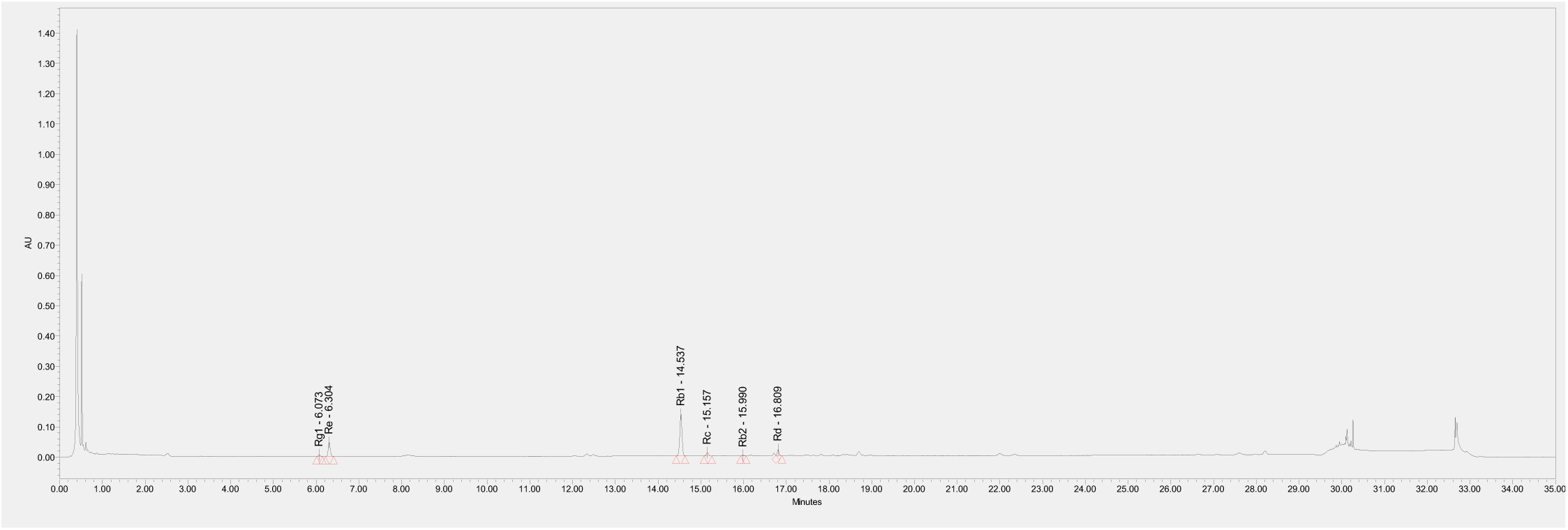
Typical chromatograms of American ginseng extract by HPLC-UV.

**Supplementary Figure 2.**
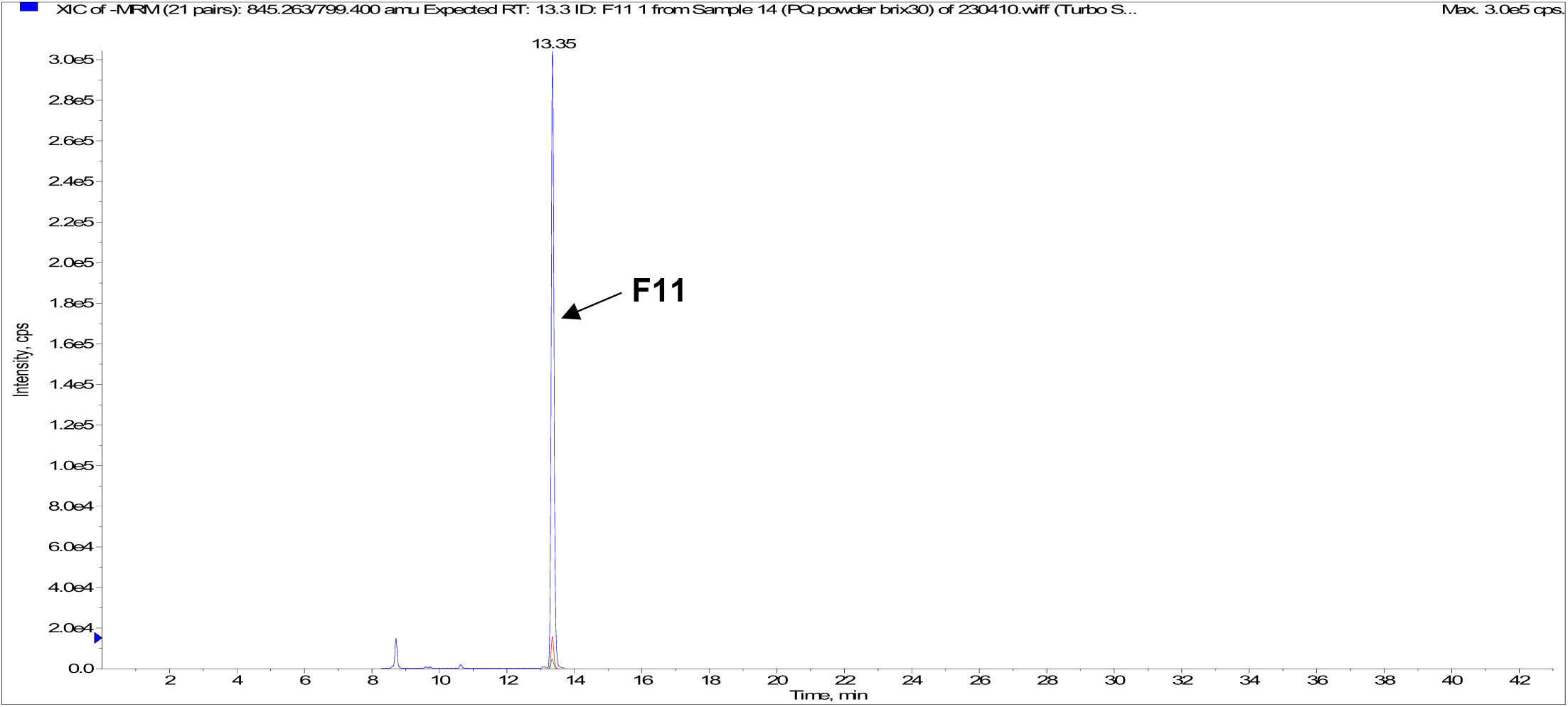
Typical chromatograms of American ginseng extract by UHPLC-MS/MS.

## Notes

Conflict of interest statement: Jaehoon Lee and Byung-Cheol Han are employees of Korea Ginseng Corporation. Other authors declare no competing financial interests.

### Competing Interest Statement

Jaehoon Lee and Byung-Cheol Han are employees of Korea Ginseng Corporation. Other authors declare no competing financial interests.

